# Polygenic mutations model the pleiotropic disease of Fanconi Anemia

**DOI:** 10.1101/2020.09.01.277038

**Authors:** Karl-Heinz Tomaszowski, Sunetra Roy, Caezaan Keshvani, Martina Ott, Courtney DiNardo, Detlev Schindler, Katharina Schlacher

## Abstract

Fanconi Anemia (FA) is a prototypic genetic disease signified by heterogeneous phenotypes including cancer, bone marrow failure, short stature, congenital abnormalities, infertility, sub-mendelian birth rate, genome instability and high cellular sensitivity to cancer therapeutics^1-4^. Clinical diagnosis is confirmed by identifying biallelic, homo- or hemizygous mutations in any one of twenty-three *FANC* genes^1,5^. Puzzlingly, inactivation of one single *Fanc* gene in mice fails to faithfully model the human disease manifestations^6-8^. We here delineate a preclinical *Fanc* mouse model with mutations in two genes, *Fancd1/Brca2* and *Fanco/Rad51c*, that recapitulates the severity and heterogeneity of the human disease manifestations including death by cancer at young age. Surprisingly, these grave phenotypes cannot be explained by the sum of phenotypes seen in mice with single gene inactivation, which are unremarkable. In contrast to expectations from classic epistasis analysis of genetic pathways, the data instead reveal an unexpected functional synergism of polygenic *Fanc* mutations. Importantly in humans, whole exome sequencing uncovers that *FANC* co-mutation in addition to the identified inactivating *FANC* gene mutation is a frequent event in FA patients. Collectively, the data establish a concept of polygenic stress as an important contributor to disease manifestations, with implications for molecular diagnostics.

## Main

Homozygous knock-outs or mutations in any one *FANC* gene alone reported so far do not cause FA in mice but instead only mild and selective disease phenotypes^6-8^. We sought to genetically test if combining *FANC* mutations could provide an improved mouse model that better reflects the variety of phenotypes seen in humans. Several FANC genes are essential for embryonic survival, including *FANCD1/BRCA2* and *FANCO/RAD51C*, and their gene knock-out (KO) is not viable in humans or mice^9,10^. FA patient mutations on the other hand frequently are small base pair-level pathogenic variants^11-13^. We created a viable homozygous *Fanco/Rad51c* mutant mouse using CRISPR/Cas9 technology carrying a six base-pair deletion mutation resulting in a deletion of two leucines at positions 270/271, corresponding to human L261/L262 (*Rad51c*^*ΔL270-_L271/ΔL270_L271*^, designated *Rad51c*^*mut*^ for the homozygous mutant mice from here on; Fig. 1a, b, and Extended Data Fig. 1a-c). The deletion is located within an *α*-helix shared with the reported FANCO mutation R258H^13^, corresponding to murine R267 (Extended Data Fig. 1d). A two amino-acid deletion within an *α*-helix causes a rotation of the helix and thus affects the location, and so perhaps also function, of the arginine critical to suppress disease. In contrast to *Rad51c* KO mice, the mutant mice express the truncated protein (Fig. 1c), supporting their viability.

**Fig. 1.**
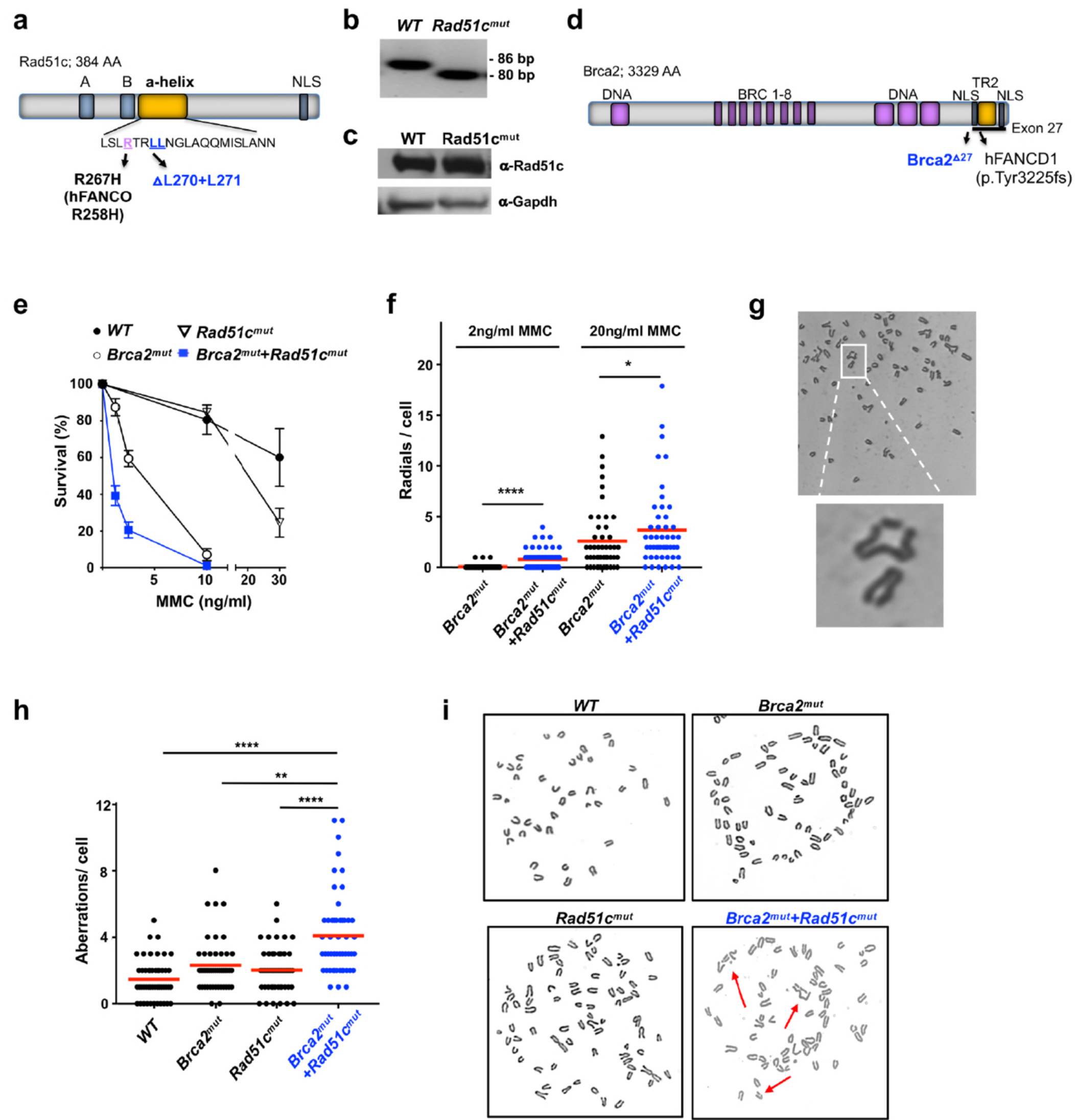
Polygenic *Brca2+Rad51c* mutations confer hyper-genome instability. **a**, Schematic of murine Rad51c. Depicted is the *α*-helix that contains the RAD51C/FANCO mutation identified in the FANCO patient (human R258H, murine R268), and the adjacent site for the two-amino acid deletion in the *Rad51c*^*dL270,L271/dL270,L271*^ (*Rad51c*^*mut*^) mouse. AA, amino acids; A,B, Walker A and B motifs; NLS, nuclear localization sequence. **b**, PCR genotyping of ear biopsies showing the six base pair (bp) deletion. **c**, Western blot of wild-type (WT) and *Rad51c*^*mut*^ mouse testis cell extracts showing similar protein expression. **d**, Schematic of murine Brca2. Depicted are the BRCA2/FANCD1 mutation identified in the FANCD1 patient (c.9672dupA), which results in a truncation of exon 27 (p.Tyr3225fs*30). The *Brca2*^*Δ27/Δ27*^ (*Brca2*^*mut*^) mouse bears an exon 27 deletion likewise truncating *Brca2* similar to the human FANCD1 truncation. AA, aminoacids; BRC, Rad51 binding motifs; TR2, Rad51 stabilization domain; DNA, DNA binding folds/motifs; NLS, nuclear localization sequence. **e**, Cellular sensitivity of bone marrow cells to varying concentrations of Mitomycin C (MMC) as indicated. The number of colonies of progenitor cells were determined after 14 days. **f**, Scatter dot blot of radial chromosome structures in metaphase spreads of mouse adult fibroblasts (MAF) after treatment with 2 or 20ng/ml MMC as indicated. **g**, Representative images of metaphase chromosome spreads in *Brca2*^*mut*^*+Rad51c*^*mut*^ MAFs **h**, Scatter dot blot of spontaneous chromosome aberrations in metaphase spreads in MAFs. **i**, Representative images of metaphase chromosome spreads without exogenous damage. Error bars represent the standard error of the mean. Bars represent the mean of compiled data from biological repeats. *p-*values are derived using the Anova test. * *p* < 0.05, **P < 0.01, *** *p* < 0.001, **** *p* < 0.0001.

Similar to *Rad51c, Brca2* gene KO is incompatible with cellular and organismal viability^9^. Mice with C-terminal *Brca2* gene truncation^14^ (*Brca2*^*Δ27/Δ27*^, designated *Brca2*^*mut*^ for the homozygous mutant mice from here on) that are similar to a pathogenic variant found in FANCD1 patients (BRCA2 p.Tyr3225fs*30)^12^; Fig. 1d), on the other hand are viable. Despite functional defects of the truncating mutation in particular in replication stability by fork protection^15^, the mice do not develop FA, albeit they show mildly reduced fertility and an increased risk of cancer with a late-age onset at >60 weeks^14^. As a best approximation to patient FANC mutations, we crossed the *Rad51c*^*mut*^ with *Brca2*^*mut*^ mice. Polygenic homozygous *Brca2*^*mut*^*+Rad51c*^*mut*^ mice are viable, but the frequency from polygenic heterozygous intercrosses is three-fold lower compared to the expected Mendelian birth rate (*p* = 0.02; Extended Data Table 1).

Hyper-sensitivity to DNA crosslinking agents and high genome instability are key-features of FA patient cells^3,4^. Using a colony formation assay, we see that bone marrow cells from *Brca2*^*mut*^*+Rad51c*^*mut*^ mice show severe cellular sensitivity to mitomycin C (MMC), consistent with an FA cell profile (Fig. 1e). In contrast, cells from *Rad51c*^*mut*^ or *Brca2*^*mut*^ mice only exhibit very mild to moderate sensitivity (Fig. 1e). Metaphase spreads of primary ear fibroblasts exposed to a high concentration of MMC (20 ng/ml) show increased chromosomal aberrations in both *Rad51c*^*mut*^ and *Brca2*^*mut*^ fibroblasts compared to wild-type fibroblasts, while *Brca2*^*mut*^*+Rad51c*^*mut*^ fibroblasts had significantly more aberrations than any of the other genotypes (Extended Data Fig. 2a, b). Interestingly, we observed that radial structures were less pronounced in *Brca2*^*mut*^ fibroblasts with a lower concentration of MMC (2 mg/ml), compared to what is seen in *Brca2*^*mut*^*+Rad51c*^*mut*^ fibroblasts, suggesting that the polygenic mutant mice are sensitive even at low amounts of DNA damage (Fig. 1f, g). Most strikingly, without any external genotoxic stress, neither *Rad51c*^*mut*^ nor *Brca2*^*mut*^ fibroblasts showed any more measurable aberrations compared to wild-type fibroblasts (Fig 1h, I, and Extended Data Fig. 2c). In stark contrast, spontaneous chromosomal aberrations are significantly increased in *Brca2*^*mut*^*+Rad51c*^*mut*^ fibroblasts (Fig 1h, I, and Extended Data Fig. 2c), suggesting strong intrinsic genomic vulnerability of the polygenic FANC mutant cells.

**Fig. 2.**
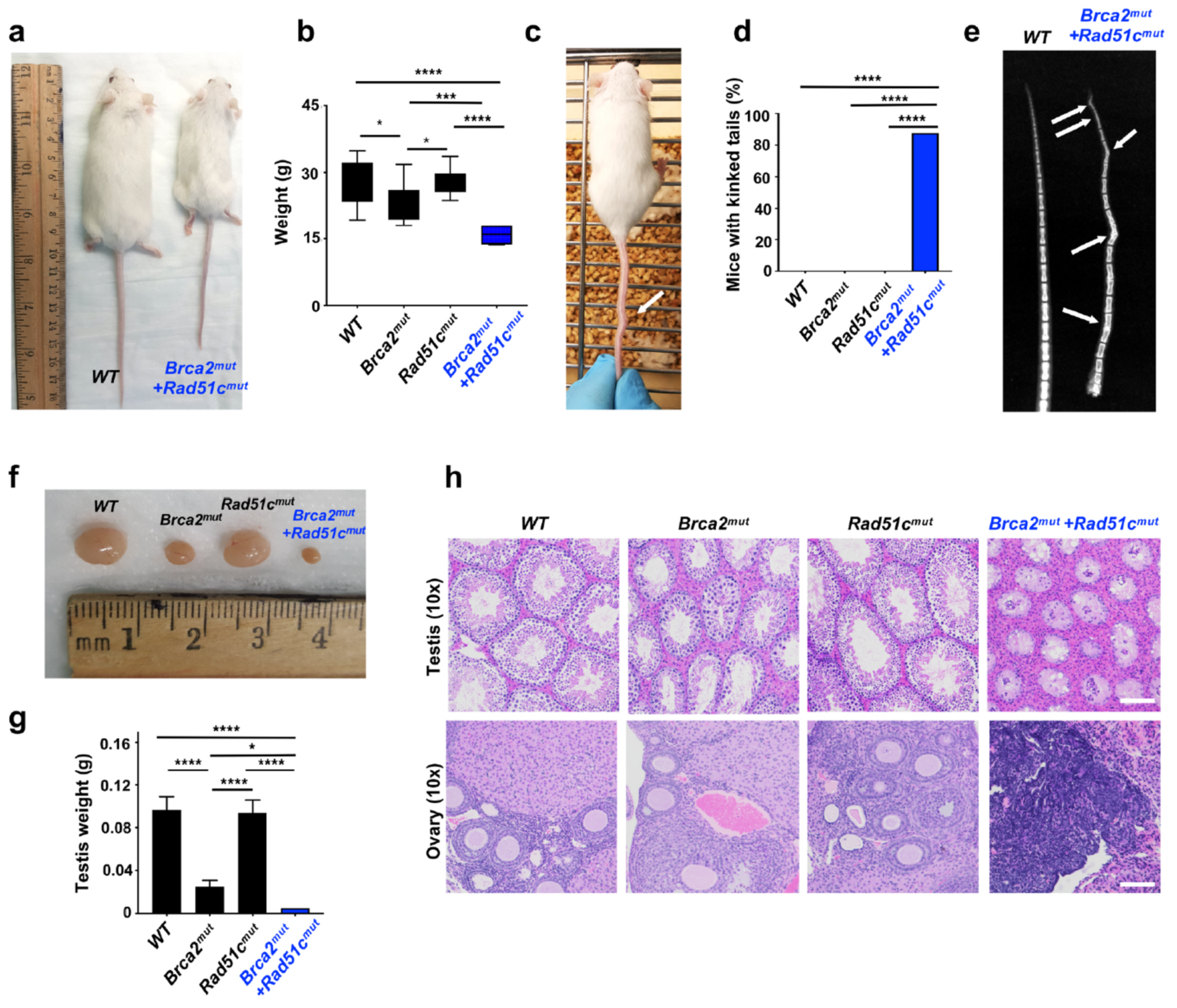
Polygenic *Brca2+Rad51c* mutant mice show congenital deformations. **a**, Representative image of wild-type (WT) and *Brca2*^*mut*^*+Rad51c*^*mut*^ mice at 12 weeks exhibiting noticeable growth defect. **b**, Box-whisker plot of weight of mice with indicated genotypes at 12 weeks (n=7-20 mice per group). **c**, Representative image of *Brca2*^*mut*^*+Rad51c*^*mut*^ mouse with kinked tail (arrow). **d**, Box plot of frequency of mice with kinked tails with indicated genotypes (n=10 mice). **e**, Representative X-ray image of tails of WT and *Brca2*^*mut*^*+Rad51c*^*mut*^ mice indicating skeletal abnormalities (arrows). **f**, Representative image of testis of mice with indicated genotypes at 12 weeks. **g**, Bar graph of weight of testis with indicated genotypes at 12 weeks. **h**, Representative images of hematoxylin and eosin (H&E)-stained tissue cross-sections of testis and ovary, respectively, from animals with indicated genotypes at 12 weeks. Scale bars indicate 100μm. Error bars represent the standard error of the mean. *p-*values are derived using the ANOVA test. * *p* < 0.05, *** *p* < 0.001, **** *p* < 0.0001.

FA patients frequently are born with congenital abnormalities causing small stature and skeletal malformations^1,5^. *Brca2*^*mut*^*+Rad51c*^*mut*^ mice have a significantly reduced weight and are small, which in contrast to the monogenic mutant mice is readily discernable from an early age (Figs. 2a, b). Additionally, *Brca2*^*mut*^*+Rad51c*^*mut*^ mice, but not single mutant *Rad51c*^*mut*^ or *Brca2*^*mut*^ mice, exhibit visible skeletal deformations with kinked tails (Fig. 2c, d). We further examined the tails by X-ray imaging to determine the cause of the tail kinks and found that many of the tail vertebrae in *Brca2*^*mut*^*+Rad51c*^*mut*^ mice are fused and exhibit an abnormal, thickened vertebrae shape (Fig. 2e).

Congenital abnormalities can cause infertility, often observed in FA patients^16^. We examined the primary sex organs in the mice and found that while the testis of *Brca2*^*mut*^ mice are smaller compared to *Rad51c*^*mut*^ mice or wild-type mice, the testis size in mice with polygenic *Brca2*^*mut*^*+Rad51c*^*mut*^ mutations are further significantly reduced (Fig. 2f, g). Histological examination of testis cross-sections using Hematoxylin and Eosin staining shows that the seminiferous tubuli in *Rad51c*^*mut*^ mice testis appear normal, while *Brca2*^*mut*^ mice show a greater heterogeneity accompanied with some expressing a mild decrease in spermatocytes residing in the lumen of the tubuli (Fig. 2h and Extended Data Fig. 3a), consistent with the mild fertility defect previously reported in this mouse model^14^. In stark contrast, the polygenic *Brca2*^*mut*^*+Rad51c*^*mut*^ mice shows a complete loss of spermatogonia germ cells in addition to all spermatocytes (Fig. 2h, upper panels, and Extended Data Fig. 3a). The data uncovers a Sertoli-cell only phenotype in these mice, which is a common FA phenotype^16^ and causes complete infertility. Similarly, cross-sections of ovaries show the complete loss of ovarian follicles in the *Brca2*^*mut*^*+Rad51c*^*mut*^ mice, but not in *Rad51c*^*mut*^, *Brca2*^*mut*^, or wild-type mice (Fig. 2h, lower panels, and Extended Data Fig. 3b, c). Collectively, the polygenic but not monogenic mutant mice show severe birth defects resembling those seen in FA patients.

**Fig. 3.**
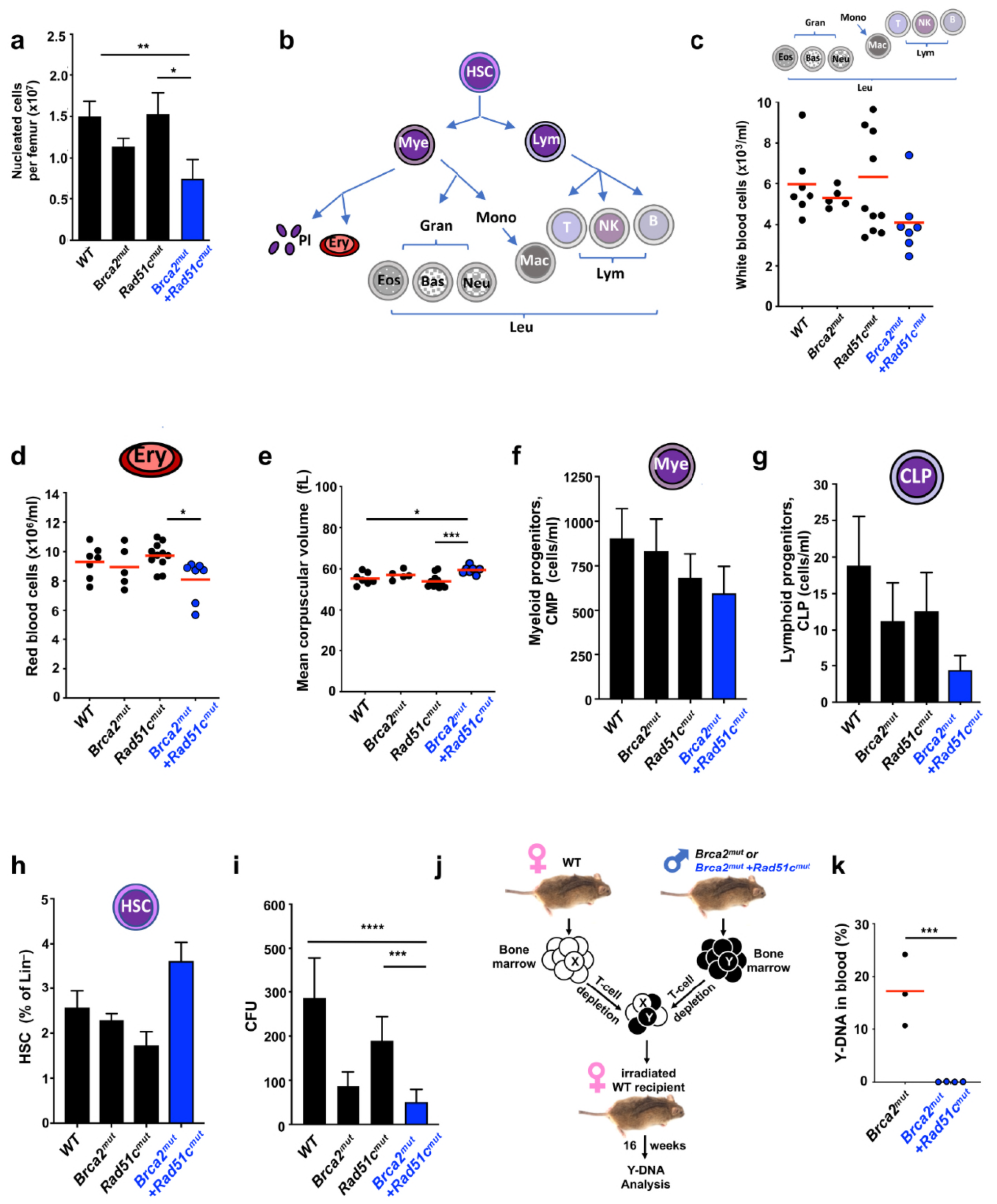
Hematological analysis of young polygenic *Brca2+Rad51c* mutant mice reveals pre-state of bone-marrow failure. **a**, Bar graph of nucleated cells per femurs of mice between 8-12 weeks (*n* = 3-7). **b**, Schematic of hematological cell lineages. HSC, hematopoietic stem cell. Mye, myeloid progenitor. CLP, common lymphoid progenitor. Pl, platelets. Ery, Erythrocytes, red blood cells. Leu, Leukocytes, white blood cells. Gran, Granulocytes. Eos, Eosinophils. Bas, Basophiles. Neutr, Neutrophils. Mono, Monocytes. Macro, Macrophages. Lym, Lymphocytes. T, T-cells. B, B-cells. NK, natural killer cells. **c-e**, Complete blood count analysis of 8–12 weeks old *Brca2*^*mut*^*+Rad51c*^*mut*^ and control mice (*n* = 6-12). (**C**) Peripheral white cell concentration. (**D**) Peripheral red cell concentration. (**E**) Mean corpsular volume as a measure for macrocitosis. **f-i**, Hematological lineage analysis by FACS was performed for bone marrow cells collected from 8-12 weeks old mice with indicated genotypes (n=5-6). **f**, Bar graph of common myeloid progenitor population **g**, Bar graph of common lymphoid progenitor population. **h**, Bar graph of hematopoietic stem cell population. **i**, Bar graph of colony formation capacity (CFU) of bone marrow progenitor cells per 30.000 plated cells. **j**, Schematic of 16-week repopulation assay work flow. Briefly, bone marrow cells from female wild-type (WT) and male mutant mice were mixed and allowed to repopulate in female mice that were depleted for their bone-marrow. After 16 weeks, the repopulation capacity of mutant bone marrow is measured by PCR of male cells in the blood, which were contributed by the mutant mouse. **k**, Quantitation of repopulation assay. Error bars represent the standard error of the mean. Red bar in scatter graphs represent the mean. *p-*values (A-I) are derived using the ANOVA test. * *p* < 0.05, *** *p* < 0.001, **** *p* < 0.0001. The *p-*values (K) is derived using the Student T-test. *** *p* < 0.001.

FA most frequently leads to bone marrow failure or aplastic anemia, characterized by a hypocellular bone marrow with progressive peripheral pancytopenia resulting in low blood cell counts. We first counted the number of bone marrow cells, which are lowest in *Brca2*^*mut*^*+Rad51c*^*mut*^ mice (Fig. 3a). To further test if any blood cells are preferentially affected, we performed a standard blood cell analysis by complete blood count on young mice (age 8-12 weeks, Fig. 3b-e and Extended Data Fig. 4a-h). While the number of platelets appears similar in all genotypes at this age (Fig. S4A), there is a small but consistent decrease in the white blood cells in the *Brca2*^*mut*^*+Rad51c*^*mut*^ mice compared to the single-mutant mice (Fig. 3c, d). More specifically, differentiated cells from both myeloid and lymphoid linages are reduced in the polygenic mice, including monocytes, granulocytes and leucocytes (Extended Data Fig. 4b-f). In addition, the red blood cell counts are significantly reduced in the double-mutated mice compared to *Rad51c*^*mut*^ mice. (Fig. 3d). Early aplastic anemia is accompanied by macrocytosis, which is one of the earliest hematological disease indicators in FA^17^. Consistent with the manifestations in FA patients, *Brca2*^*mut*^*+Rad51c*^*mut*^ mice but not *Rad51c*^*mut*^ nor *Brca2*^*mut*^ mice show significantly enlarged red blood cells, suggesting macrocytosis and abnormal hematopoiesis (Fig. 3f, and Extended Data Fig. 4g, h). Collectively, the data obtained from differentiated cells of myeloid and lymphatic lineages in young polygenic *Brca2*^*mut*^*+Rad51c*^*mut*^ mutant mice is consistent with early bone marrow failure.

**Fig. 4.**
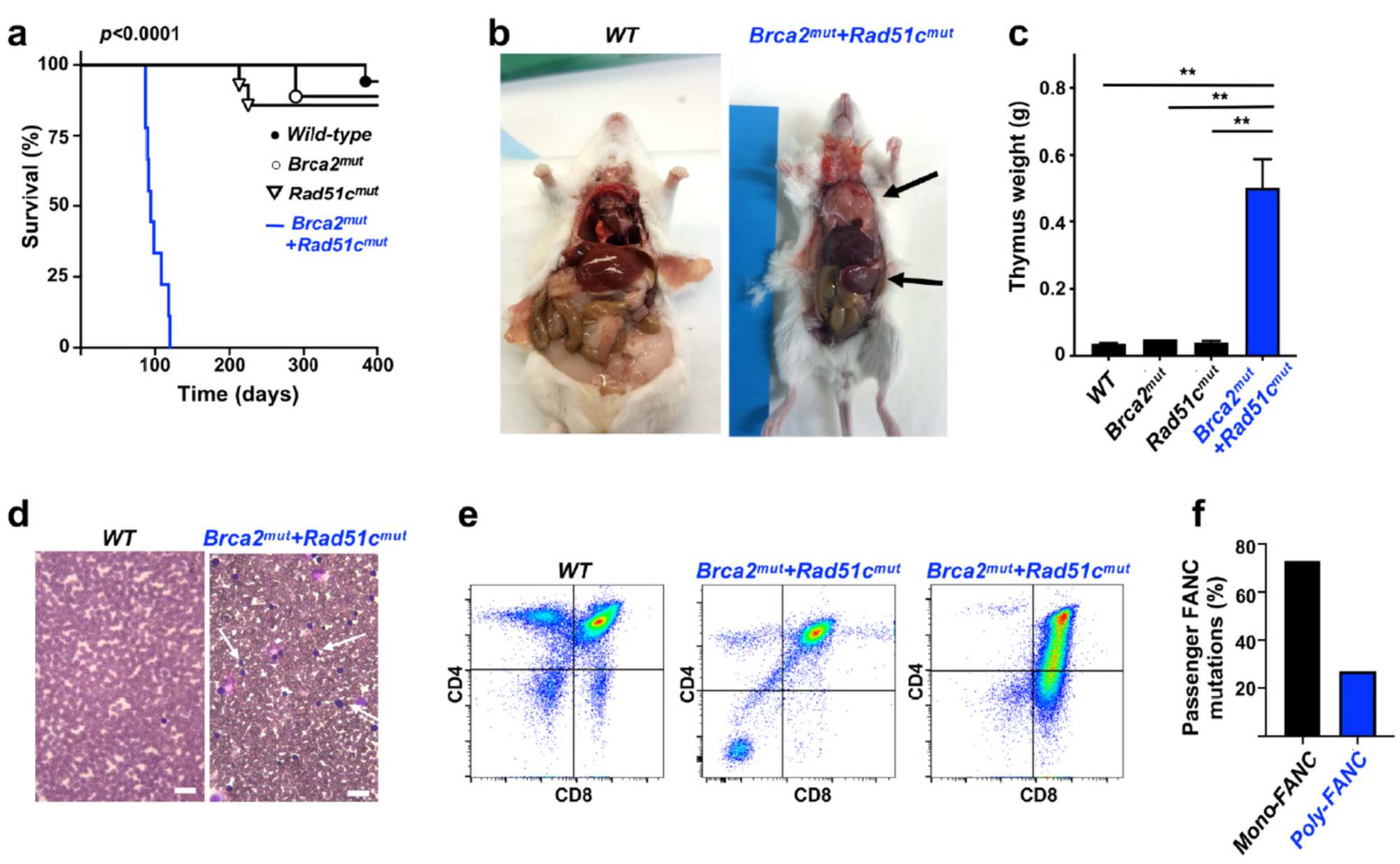
Polygenic *Brca2+Rad51c* mutant mice develop cancer. **a**, Kaplan-Meier curves for overall survival of wild-type (WT), *Brca2*^*mut*^, *Rad51c*^*mut*^, and *Brca2*^*mut*^*+Rad51c*^*mut*^ mice. n=9-17 per genotype. *Brca2*^*mut*^*+Rad51c*^*mut*^ against all other genotypes: p<0.0001; WT against *Brca2*^*mut*^: *p*=0.9890; WT against *Rad51c*^*mut*^: *p*=0.4957; *Brca2*^*mut*^ against *Rad51c*^*mut*^: *p*=0.8058. **b**, Representative images of malade *Brca2*^*mut*^*+Rad51c*^*mut*^ and age-matched WT mice. Top arrow shows large mediastinal thymic mass found in sick *Brca2*^*mut*^*+Rad51c*^*mut*^ mice Lower arrow points to enlarged spleen. **c**, Bar-graph of thymus mass quantitation in malade *Brca2*^*mut*^*+Rad51c*^*mut*^ and age-matched control mice (n=3-5). **d**, Representative images of blood films (40x magnification) obtained from malade *Brca2*^*mut*^*+Rad51c*^*mut*^ mice and age-matched WT mice shows high number of abnormal leukocytes (arrow). Scale bar, 30μm. **e**, FACS analysis of thymus cells with antibodies against CD4 and CD8. Cells from malade *Brca2*^*mut*^*+Rad51c*^*mut*^ mice (middle and right) show clonal expansion of early T-cells, indicating the development of acute lymphoblastic leukemia. **f**, Exome sequencing of 37 confirmed FA patient serum filtered for additional passenger mutations defined as non-synonymous FANC mutations in additional FANC genes other than the primary, diagnostic biallelic FANC mutation. Error bars represent the standard error of the mean. *p-*values (C) are derived using the ANOVA test. ** *p* < 0.01, *p-*values.

To further trace the cells of origin for the hematological defect, we performed hematopoietic lineage FACS analysis^18^. Consistent with the reduced counts in differentiated cells, the bone marrow of *Brca2*^*mut*^*+Rad51c*^*mut*^ mice contain less of both, myeloid and common lymphoid progenitor cells, albeit the numbers do not allow for statistical significance in the young mice (Fig. 3f, g, and Extended Data Fig. 5a, b). Intriguingly, the frequency of the hematopoietic stem cells (HSC) on the other hand is slightly increased in polygenic compared to single mutant mice at the time-point measured (Fig. 3h). Increased HSCs are a sign of stress-induced HSC proliferation, that can lead to their depletion^19^. We therefore sought to further test bone marrow cell functions. Colony forming unit assays revealed a functional defect of *Brca2*^*mut*^*+Rad51c*^*mut*^ bone marrow cells, albeit *Brca2*^*mut*^ proliferation-potent cells too show a clear reduction in colony formation capacity (Fig. 3i). To stringently test HSC function, we performed a bone marrow repopulation assay by injecting an equal amount of bone marrow cells from of either male *Brca2*^*mut*^*+Rad51c*^*mut*^ and female wild-type mice, or male *Brca2*^*mut*^ and female wild-type mice into irradiated mice that are so bone marrow-depleted (Fig. 3j, k). Strikingly, quantifying the frequency of Y-chromosome DNA in blood cells 16 weeks after transplantation we found that while bone marrow of *Brca2*^*mut*^ mice is significantly contributing to the make-up of the blood cells in the experimental animal, *Brca2*^*mut*^*+Rad51c*^*mut*^ bone marrow cells completely fail to do so (Fig. 3k). Thus, the data demonstrates a dramatic loss of HSC function *in vivo* in *Brca2*^*mut*^*+Rad51c*^*mut*^ bone marrow cells.

We measured the overall survival of the genetic mouse model. Dramatically, all *Brca2*^*mut*^*+Rad51c*^*mut*^ mice die between 3-4 months of age (87-120 days, Fig. 4a). This is in stark contrast to wild-type and either of the single-gene mutant mice, where neither *Rad51c*^*mut*^ nor the *Brca2*^*mut*^ mice show a significant difference in overall survival in the first year of life (Fig. 4a).

Patients with Fanconi anemia related to loss of FANCD1/BRCA2 exhibit a variety of cancers, including brain tumors, Wilms tumors, acute myeoloid leukemia and T-cell leukemia^3,11,20^. We found that the polygenic mutant mice have a strikingly enlarged thymus and spleen (Fig. 4b, c and Extended Data Fig. 6a), suggestive of hematologic malignancy. Consistently, thymus cells of polygenic mutant mice are greatly enlarged in size, indicating active proliferation (Extended Data Fig. 6b). Moreover, leukemic mice have an unusually elevated number of white blood cells (Extended Data Fig. 6c), and blood smears reveal homogenous appearing blast cells (Fig. 4d, white arrow), collectively indicating the presence of leukemia. Additional hematopoietic lineage analysis by FACS determined strong abnormal T-cell differentiation of thymus cells in *Brca2*^*mut*^*+Rad51c*^*mut*^ mice (Fig. 4e). Together, the analysis suggests that the polygenic *Brca2*^*mut*^*+Rad51c*^*mut*^ mice die of T-cell leukemia, consistent with the FANCD1 patient phenotypes.

To test if polygenic FANC inactivation is a general feature found in FA patients, we performed a literature search and found multiple examples of FA patients with *FANC* gene mutations in addition to the *FANC* inactivating variants that the patient’s disease is attributed to (Extended Data Table 2). To rigorously test for polygenic FANC mutations in FA patients, we performed whole-exome sequencing on DNAs of 37 FA patients with known biallelic mutations in one given FANC gene. Strikingly, 27% of the patients tested harbor non-synonymous germline mutations in additional FANC genes (Fig. 4f), suggesting a high prevalence of polygenic FANC mutations in patients.

Clinically, FA is defined as a biallelic inactivation in one of 23 so far identified FANC genes^1,5^. Perhaps forgotten, Guido Fanconi, the Swiss pediatrician who the disease is named after, speculated that the disease was too complex to be caused by one gene^21^. We here show that in contrast to monogenic *Fanc* mutations, polygenic *Fanc* gene mutations in mice closely recapitulates FA disease manifestations found in patients, providing a comprehensive preclinical mouse model of FA. While the mice succumb to cancer before developing severe bone marrow failure, the hematological analysis of young polygenic mutant mice revealed macrocytosis, which in patients is indicative of hematopoietic stem cell depletion leading to aplastic anemia and amongst others can be caused by erythropoietic stress and DNA replication defects^17,22^. Interestingly, externally applied agents that cause DNA damage and DNA replication stress by aldehydes^23^ or DNA crosslinks^24,25^ in mono-genic *FANC* mutant mice significantly worsens the mice’s phenotypes and better model the human FA disease manifestations compared to the genetic alterations alone. We therefore suggest that the genetic combination of FANC inactivation could cause endogenous stress, in analogy to the principle of oncogene induced replication stress^26^. In support of this idea, cells from polygenic mice exhibit significantly higher spontaneous chromosomal aberrations in the absence of exogenous DNA damage. Our proposed concept of “genetic stress” by compounding genetic alterations is also affirmed by the observation that the same FANC gene knockouts causes varying phenotypic severity depending on the genetic background of the pure-bred animals^7,27,28^, supporting the concept that the genetic background can shape the disease development.

FA occurs with a frequency of 1:100,000-250,000. Inconsistently, ∼4.3% of individuals of the Western population carry heterozygous disease-causing FANC variants, making the disease far less frequent than what would be expected from Mendelian transmission^29,30^. The requirement for other factors in addition to biallelic FANC mutations in one gene at least in part could account for this discrepancy. Such factors could include additional genetic stress, or separately additional physical stress by DNA damage such as from aldehydes. Alternatively, the additional synergistic mutations may not be restricted to so far identified FANC genes but feasibly can include many of the DNA repair pathway proteins that FANC genes are known to interact with. Moreover, aldehyde stress has been shown to cause BRCA2 protein destabilization^31^. Thus, externally triggered DNA damage in principle may contribute to inducible FANC protein decreases and so effective additional FANC inactivation without genetic alterations. The data here show that polygenic FANC mutations are a frequent event in FA patients lending support to the concept of polygenic stress as a disease inducing mechanism. It further suggests that additional gene analysis could aid in predicting the severity of the disease as an added parameter, which is of particular importance in families with heterogeneously affected siblings^32^.

The concept of requiring polygenic gene inactivation for disease manifestation may usefully inform other rare genetic disease models beyond FA. In a so far experimentally untested mathematical model, the co-recessive inheritance theory by Lambert and Lambert predicted that diseases caused by mutations in other DNA damage response genes may require more than one gene inactivation^33^, including ataxia telengiectasia (AT associated with mutations in ATM), xeroderma pigmentosum and Cockayne syndrome (XP and CS involving nucleotide excision repair genes), and cancer. A requirement of polygenic stress by synergistic gene mutations may at least in part resolve why single-gene inactivation in mice alone only poorly models AT and CS with mild symptoms, respectively, while additional gene inactivation results in better disease modeling^34,35^. This phenomenon was so far was unexplained. Thus, polygenic stress involvement defining disease presentation, as postulated based on the observations of our mouse model, may point to a general concept.

Notably, polygenic risk evaluation, whereby the risk scores of multiple genes are added up to a compounded risk factor, is emerging as a promising tool in predicating many rare genetic diseases, including neurodevelopmental disorders, autism, Alzheimer disease, and breast cancer^36-39^. Yet challenges remain in defining proper predictive parameters. FA cells show high genome instability, so the heightened incidence of secondary mutations in other FANC genes may not come to a surprise. In the context of classic epistasis analysis of molecular pathways, the presented mouse model however reveals an entirely unexpected functional synergism of genetic interactions within the FA pathway that shape the disease manifestation. This predicts that the effect of synergistic mutations will not be random. The promising but so far incomplete success of polygenic risk prediction therefore merits further investigation into specific gene synergism in other diseases, including cancer. The data here therefore has additional implications for tumor passenger mutation., whereby a seemingly unremarkable mutation in one gene is asymptotic and tolerated, but within the context of an additional acquired mild somatic mutation causes severe and aggressive cancer. Collectively, the presented mouse model offers a comprehensive preclinical model to faithfully investigate diverse FA disease manifestations, and establishes the concept of polygenic stress driving disease severity as a testable principle in cancer and diverse genetic diseases.

## Methods

### Mouse models

Rad51c mutant mice was generated by pronuclear injection using the CRISPR/Cas9 genome editing system. Cas9 and sgRNA 5’-CTTCGTACTCGATTACTAAATGG-3’ were injected into pronuclear stage mouse embryos from FVB/J mice to generate founder animals. *Brca2*^*mut*^ mice, carrying a C-terminal truncation, were obtained from NCI Mouse Repository (SWR.129P2(Cg)-*Brca2*^*tm1Kamc*^/Nci, Strain code: 01XG9, SWR.129P2). Mice carrying both the *Brca2* C-terminal deletion and the *Rad51c* 6 base pair deletion were generated by crossing *Brca2* and *Rad51c* heterozygous mutant mice. The obtained offspring were subsequently intercrossed to obtain double-mutant mice and control genotypes in FVB/J SWR.129 hybrid background. Mice were maintained in pathogen-free conditions. Animal experiments were performed in accordance with approved animal protocol by the Institutional Animal Care and Use Committee of the University of Texas M. D. Anderson Cancer Center.

For genotyping, ears of mice were clipped and genomic DNA was extracted using DirectPCR Lysis Reagent (Viagen Biotech). PCR amplification was performed using GoTaq Green Master Mix (Promega) or Phusion High-Fidelity DNA polymerase (NEB) according to the manufacturer’s introduction. PCR products (Rad51c-Forward, 5’-CGTCATGACCTTGAAGATC-3’, Rad51c-Reverse, 5’-GATTATTTGCAAGGCTGATC-3’; Brca2-Forward, 5’-GAGAGCCCCATGCAGCCTCCACTTGCTGTG-3’ and Brca2-Reverse, 5’-CTGCCTCCAGAGACCTGAGCCGTC-3’) were directly analyzed by agarose gel based on product size.

### Generation of mouse adult fibroblasts (MAF)

Primary ear fibroblasts were derived from age-matched male and females. Briefly, a portion of ear was cut off, rinsed two times with PBS containing kanamycin (100 μg/mL) and digested with collagenase D/dispase II protease (4 mg/mL, respectively) for 45min at 37°C. After dilution with five times Dulbecco’s modified eagle’s high-glucose media (DMEM) containing 10% fetal bovine serum (FBS) and 5% Antibiotic-Antimycotic, cells were incubated overnight at 37°C. The following day, cells were passed through a 0.7μM cell strainer, washed with PBS and plated for cultivation standard media consist of DMEM supplemented with 10% FBS and 100units/ml Pen-Strep. MAFs were grown at 37°C and 5% CO2, routinely splitted two times per week and passages 2 to 10 were used for experiments.

### Cytogenetic Analysis

For chromosomal aberration analysis, 5 × 10^4^ mouse adult fibroblasts were plated for 24h in standard DMEM media. On the next day, exponentially growing cells were treated with indicated concentration of mitomycin C for 15 h followed by incubation with colcemid (0.1mg/ml, Gibco) for 5h. Afterwards, cells were swollen with 0.04 % KCl solution (12 min, 37°C), fixed in methanol/acetic acid (3:1), dropped onto microscope slides, stained with 5 % Giemsa solution and directly imaged with a Nikon Eclipse Ti-U inverted microscope. Images were analyzed using ImageJ software.

### Histological analysis

Testis and ovaries from adult mice were resected and fixed in 10% buffered formalin solution (Thermo Fisher Scientific) overnight and stored in 70% ethanol. Formalin-fixed tissues were paraffin embedded and tissue sections were counterstain in eosin-phloxine B solution for 30 seconds to one minute. Images were obtained using a Nikon Eclipse Ti-U inverted microscope.

### Bone marrow harvest

Bone marrow cells were isolated from femurs of 8-12 weeks old mutant mice and appropriate controls. Skin and muscle surrounding the bone was removed and bone marrow cells were isolated by flushing out cells with PBS using a 27 gauge needle. Pelleted cells were treated with red cell lysis buffer (Lonza) for two minutes to eliminate red blood cells.

### Flow cytometry

Flow cytometry was performed on freshly isolated bone marrow cells. The following antibodies were used to stain for hematopoietic stem cells (HSCs) and progenitor populations: biotin-conjugated lineage cocktail with antibodies anti-TER-119, anti-CD11b, anti-Ly-6G/Ly-6C (Gr-1), anti-CD3e and anti-CD45R/B220 (BioLegend), anti-streptavidin (APC-Cy7, BD Pharmingen) was used as secondary antibody, anti-c-kit (APC, clone 2B8, BD Pharmingen), anti-Sca-1 (PerCP-Cy5.5, clone D7, eBioscience), anti-CD34 (FITC, clone RAM34, eBioscience), anti-CD135 (Flt3) (PE, clone A2F10, eBioscience) and anti-CD127 (PE-Cy7, clone SB/199, eBioscience). For thymic T-cell maturation, cells were obtained by mashing thymus through a 0.45µM cell strainer. Afterwards, cells were stained using anti-CD3 (APC, clone 145-2C11, BD Pharmingen), anti-CD4 (PE, clone RM4-5, BD Pharmingen) and anti-CD8 (FITC, clone 53-6.7, BD Pharmingen). The samples were incubated with primary antibodies for 90 min at 4°C in the dark. Following washing steps, cells were incubated with secondary antibody for 30 min at 4°C in the dark. For quantification of cell numbers, AccuCheck counting beads (Thermo Fisher Scientific) was included in samples and calculation was performed following manufacturer’s instructions. Flow cytometric analysis was performed using a LSRFortessa X-20 Analyzer and data were analyzed with FlowJo 10.4.2 (FlowJo, LLC).

### Western blot

Testes were collected from adult male *Rad51c* mutant and wild-type mice, and protein lysates were used for immunoblotting using standard techniques.

### Peripheral blood analysis

Whole blood was collected from 8-12 weeks old mice in EDTA microvettes tubes (BD Biosciences) and complete blood count were analyzed using ADVIA 2120i Hematology systems (Siemens Healthineers) according to the manufactor’s instruction. Whole blood slides were stained with Diff-Quick stain.

### Colony formation unit assay

Hematopoietic colony formation unit assays were performed using bone marrow cells harvested as described above. The number of bone marrow cells were enumerated using trypan blue staining in a neubauer chamber (Thermo Fisher). Appropriate numbers of total bone marrow cells were then exposed to various concentration of MMC and seeded into 6-well plates with MethoCult GF M3434 (Stem Cell Technologies) media following the manufacturer’s instructions.

### X-ray imaging

X-ray imaging from mice tails was performed using an IVIS Lumina X5 imager.

### Irradiation of mice

Mice received a dose of 900 Gy of total body irradiation. Additionally, mice were treated prophylactic with enrofloxacin (Baytril) in the drinking water for one week before irradiation and for 3 weeks after irradiation.

### Competitive repopulation assay

To assess the HSC function the competitive repopulation assay was performed essentially as described previously^40^. Briefly, male bone marrow cells (5×10^4^ or 2×10^5^) from *Brca2*^*mut*^ and *Brca2*^*mut*^*+Rad51c* ^*mut*^ mice were mixed with 5×10^4^ of wild-type female bone marrow and injected into lethally irradiated (900 Gy) female wild-type recipients. Three to four recipients were used for each genotype. After 16 weeks, recipient mice were killed and genomic DNA from peripheral blood was extracted using PureLink Genomic DNA Kit (Invitrogen) according to the manufactor’s instructions. Thereafter, the relative contribution of the tested donor mutant and wild-type bone marrow to peripheral blood chimerism was assessed using quantitative PCR (qPCR) for the presence of male specific Y-chromosome (SRY-Forward, 5’-TGTTCAGCCCTACAGCCACA-3’, and SRY-Reverse, 5’-CCTCTCACCACGG-GACCAC-3’), and detection was performed with the TaqMan probe SRY-T, 5’-FAM– ACAATTGTCTAGAGAGCATGGAGGGCCA–BHQ1-3’. Primers for murine □-actin sequence were □ - actin-Forward, 5’-ACGGCCAGGTCATCACTATTG-3’, and b-actin-Reverse, 5’-ACTATGGCCTCAAGGAGTTTTGTCA-3’, and detected with the TaqMan probe □ -actin-T, 5’-FAM–AACGAGCGGTTCCGATGCCCT–BHQ1-3’. Amplification reaction was performed using TaqMan Multiplex Mix Kit (ThermoFisher) according to the manufactor’s instructions.

## Statistical analysis

Statistical data analysis was performed using GraphPad software. Unless otherwise stated, significant differences between sample groups were determined using ANOVA and statistical significance are indicated with asterisks as follows (\**p< 0*.*05*, \*\**p < 0*.*01*, \*\*\**p < 0*.*001*, \*\*\*\**p < 0*.*0001*).

### Whole exome sequencing of FA patient DNAs

Library preparation and target enrichment were achieved using the SureSelect Human All Exon system of different versions (Agilent) or TruSeq DNA Exome technology (Illumina), and was followed by next-generation sequencing on a HiSeq2500 instrument (Illumina). The average exome coverage was determined using a complete list of human exons generated by the *UCSC Table Browser*. The same procedure was performed for FA gene coverage. Generally ≥80 of the reads were on target with > 85% of bases covered at 10× depth. Data were analyzed using **Next***GENe* **Sequence Analysis software** *(Softgenetics)*. Finally, a manual filtering step based on data mining was carried out to prioritize relevant mutations. A minimum coverage by 10 reads was set as threshold for any variant to be considered faithful. The variant detection frequency was limited to a minimum of 20% of the reads covering any aberration. Potentially pathogenic variants were verified by Sanger sequencing generally using an *Applied Biosystems 3130xl* instrument.

## Data and materials availability

All data is available in the main text or the Extended data materials.

## Acknowledgments

We thank Drs. John A. Tainer, Davide Moiani, Michael Longo and Gareth Williams for sharing information and discussions on the RAD51C crystal structure, Dr. Tamara M. Haygood for advise on tail pathologies seen with X-ray, as well as the MDACC Genetically Engineered Mouse Facility (GEMF), the Small Animal Imaging Facility, the Research Histology Core Laboratory core facilities, the Advanced Cytometry & Sorting Facility at South Campus (ACSF), and the Veterinary and Comparative Pathology facility at MD Anderson Cancer Center for critical support. The work was supported by the NIEHS under award 1R01ES029680, and by CPRIT RP180463, R1312 and RP180813 (K.S.), and the FA research group at the University of Wuerzburg was supported by grants from the Schroeder Kurth Fund (D.S.). K.S. is a Rita Allen Foundation Fellow and a CPRIT scholar in Cancer Biology (previous award R1312).

## Author contributions

K.H.T. designed and performed the experiments. D.S. performed passenger mutation analysis of FA patients and a literature search. M.O., S.R., C.K. experimentally contributed to the understanding during the development of the work. C.D. contributed in the clinical understanding during the development of the work. K.S. conceived the project, advised on experimental design, performed the literature search on passenger mutation analysis, and wrote the manuscript with input from all authors.

## Additional Information

Extended Data Figures 1-5

Extended Data Tables 1-2

**Extended Data Fig. 1.**
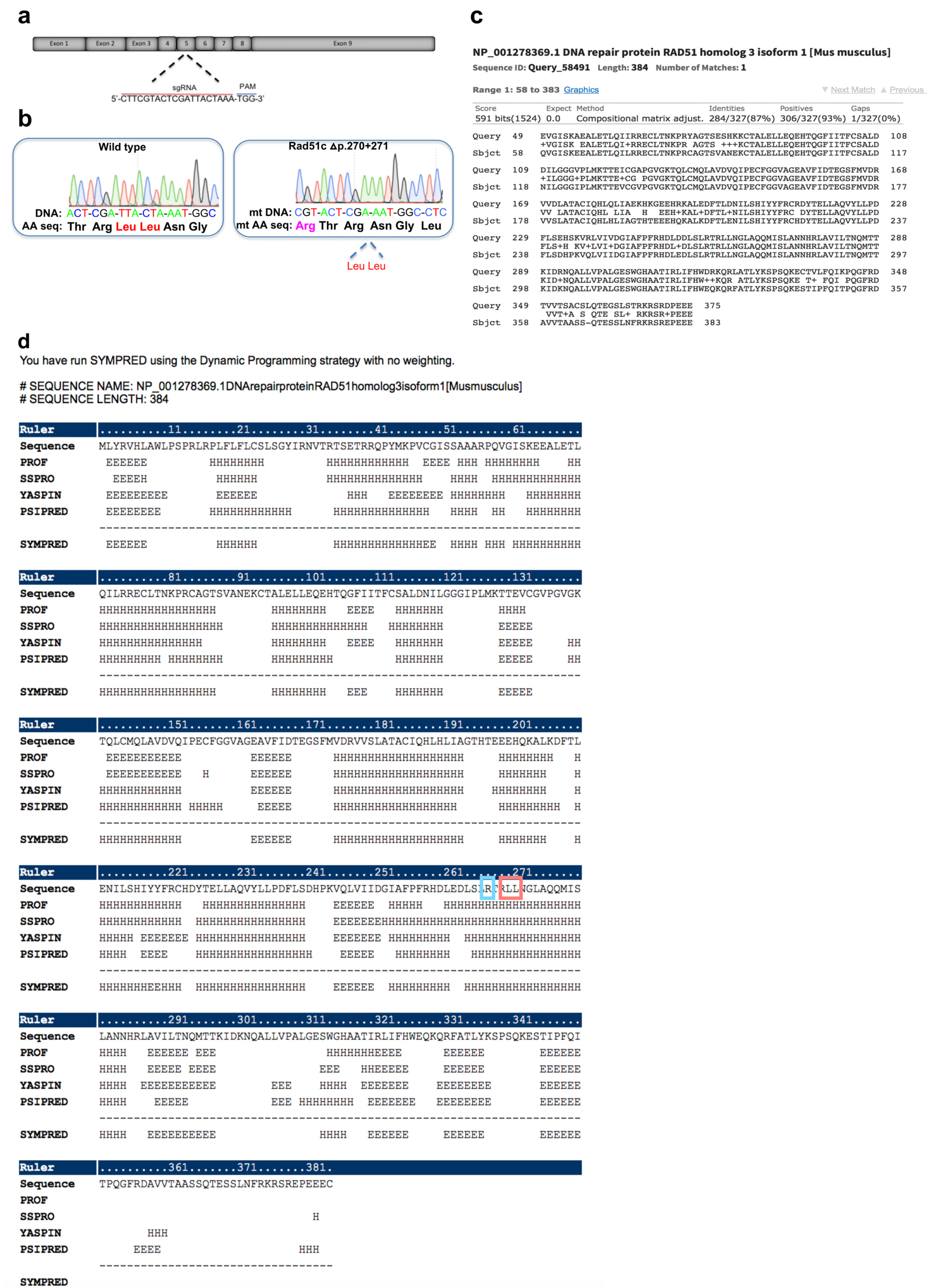
Generation of Rad51c mutant mice with 6bp deletion. a, Schematic diagram of sgRNA guide and targeting sites in the mouse *Rad51c* locus. PAM = protospacer adjacent motif. **b**, Sanger sequencing of PCR products encompassing the *Rad51c* targeting region from a representative wild-type (WT) mice and Rad51c^mut^ mouse. The two leucine residues that are deleted in Rad51c^mut^ mouse are marked in red, the adjacent Arginine that is substituted by a histidine in the FANCO patient is marked in magenta. **c**, Blast sequence alignment of human (NP_478123.1) and murine RAD51C (NP_001278369) shows high similarity and evolutionary conservation. **d**, SymPRED protein secondary structure prediction of RAD51C showing R258 (blue square, mutated to histidine in the FANCO patient) adjacent to the two leucines (red square) that are deleted in the mouse model within the same a-helix

**Extended Data Fig.2.**
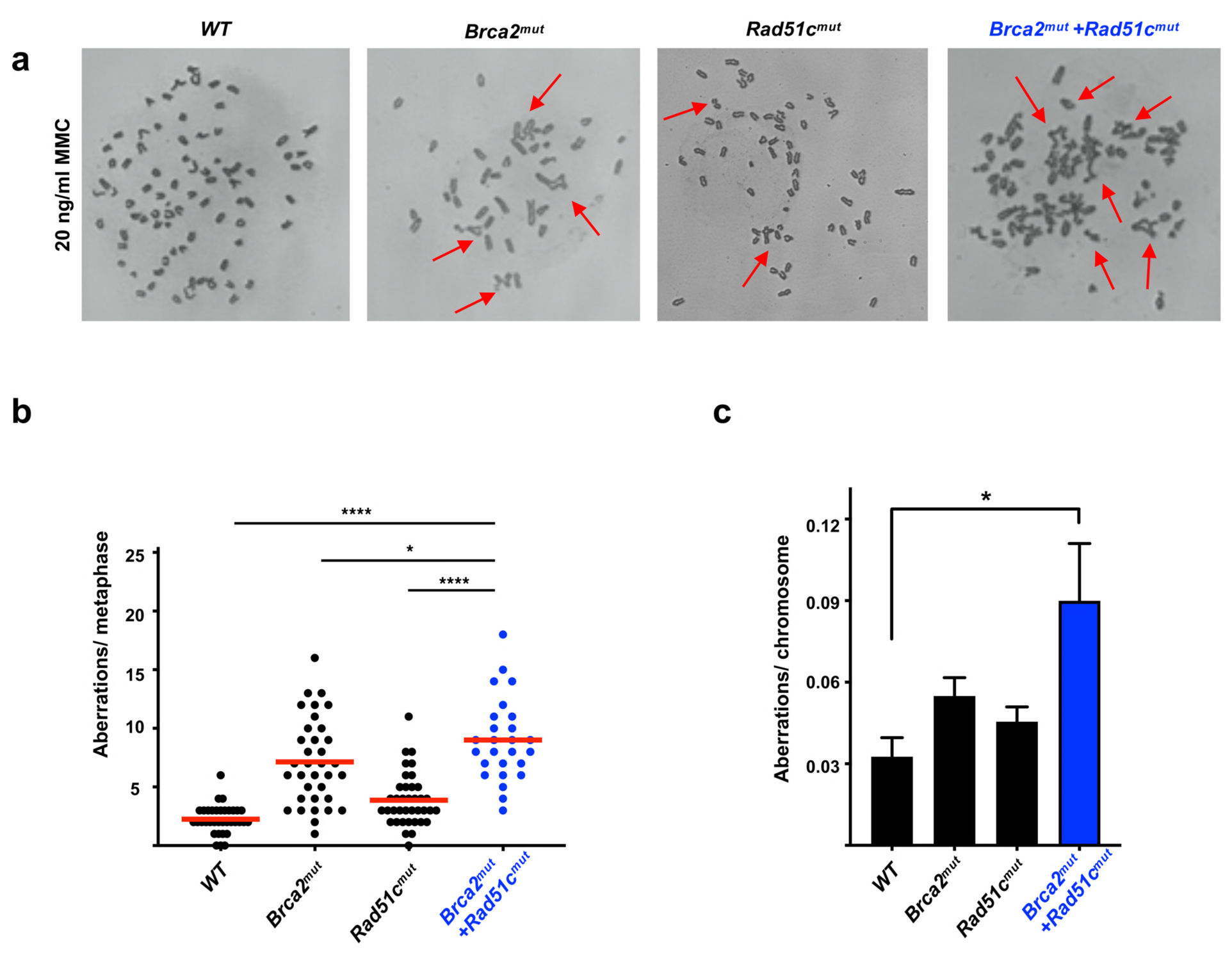
Cells with polygenic FANC mutations show increased genomic aberrations. **a**, Representative images of metaphase chromosome spreads in mouse adult fibroblasts (MAF) of indicated genotypes with 20 ng/ml mitomycin C (MMC). Examples for chromosomal aberrations are indicated with an arrow. **b**, Scatter dot blot of chromosomal aberrations in metaphase spreads of MAFs after treatment with 20ng/ml MMC as indicated. **c**, Bargraph of spontaneous chromosome aberrations per chromosome (without MMC) in metaphase spreads in MAFs. Error bars represent the standard error of the mean. The data represent the mean of compiled data from biological repeats. *p-*values are derived using the ANOVA test. * *p* < 0.05, \*\**p* < 0.01, *** *p* < 0.001, **** *p* < 0.0001

**Extended Data Fig. 3.**
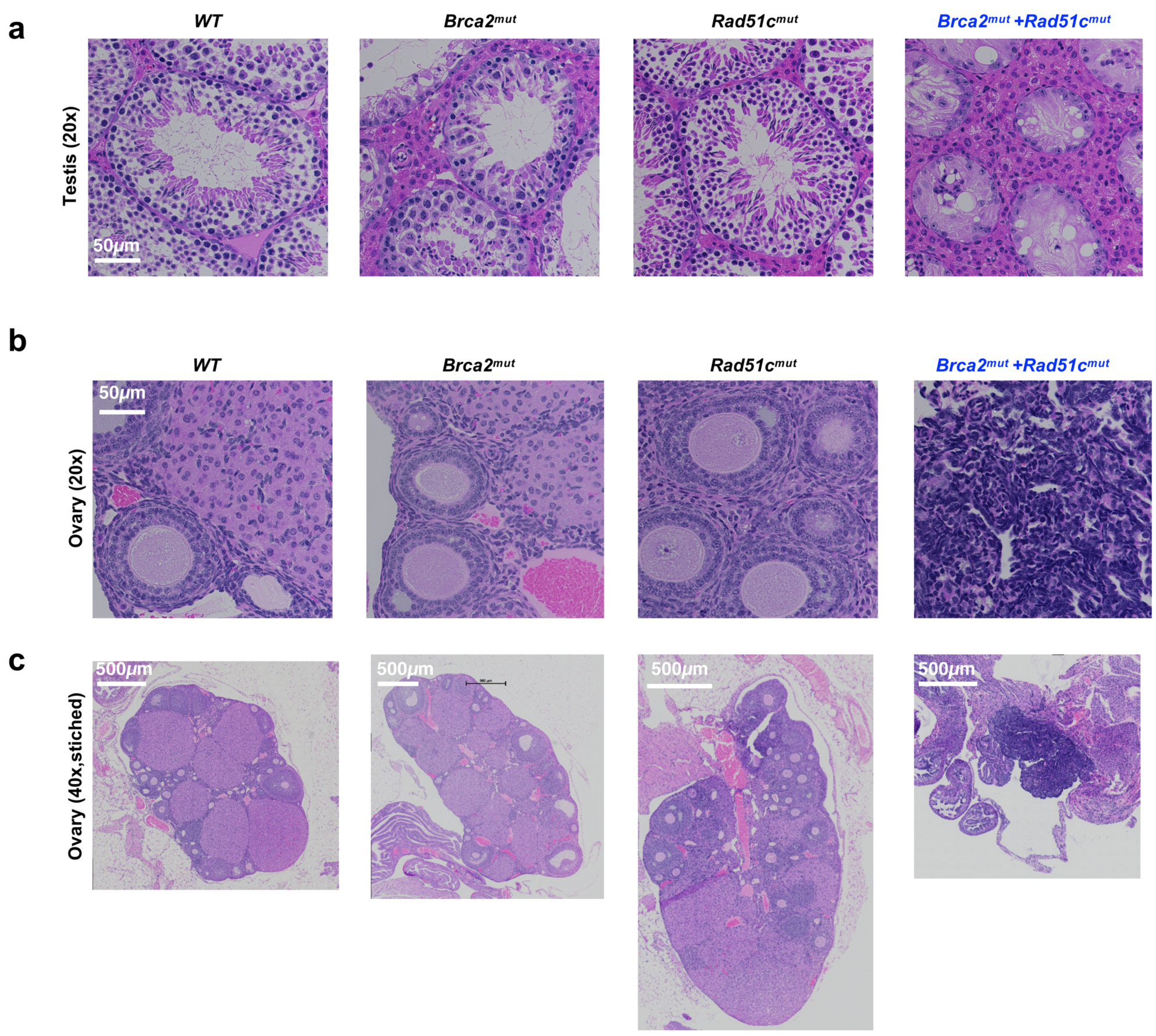
Polygenic mutant FANC mice show impaired gonad development. **a**, Representative images of hematoxylin and eosin (H&E)-stained tissue cross-sections of testis from animals with indicated genotypes at 12 weeks. Seminiferous tubuli of B*rca2*^*mut*^ mice can show decreased spermatocytes (arrow), while those of *Rad51c*^*mut*^+*Brca2*^*mut*^ mice show a Sertoli cell only phenotype with a complete loss in spermatocytes and spermatogonia. Scale bars indicate 50mm **b**, Representative images of H&E-stained tissue cross-sections of ovaries from animals with indicated genotypes at 12 weeks. Scale bars indicate 50mm **b**, and 500µm **c**.

**Extended Data Fig. 4.**
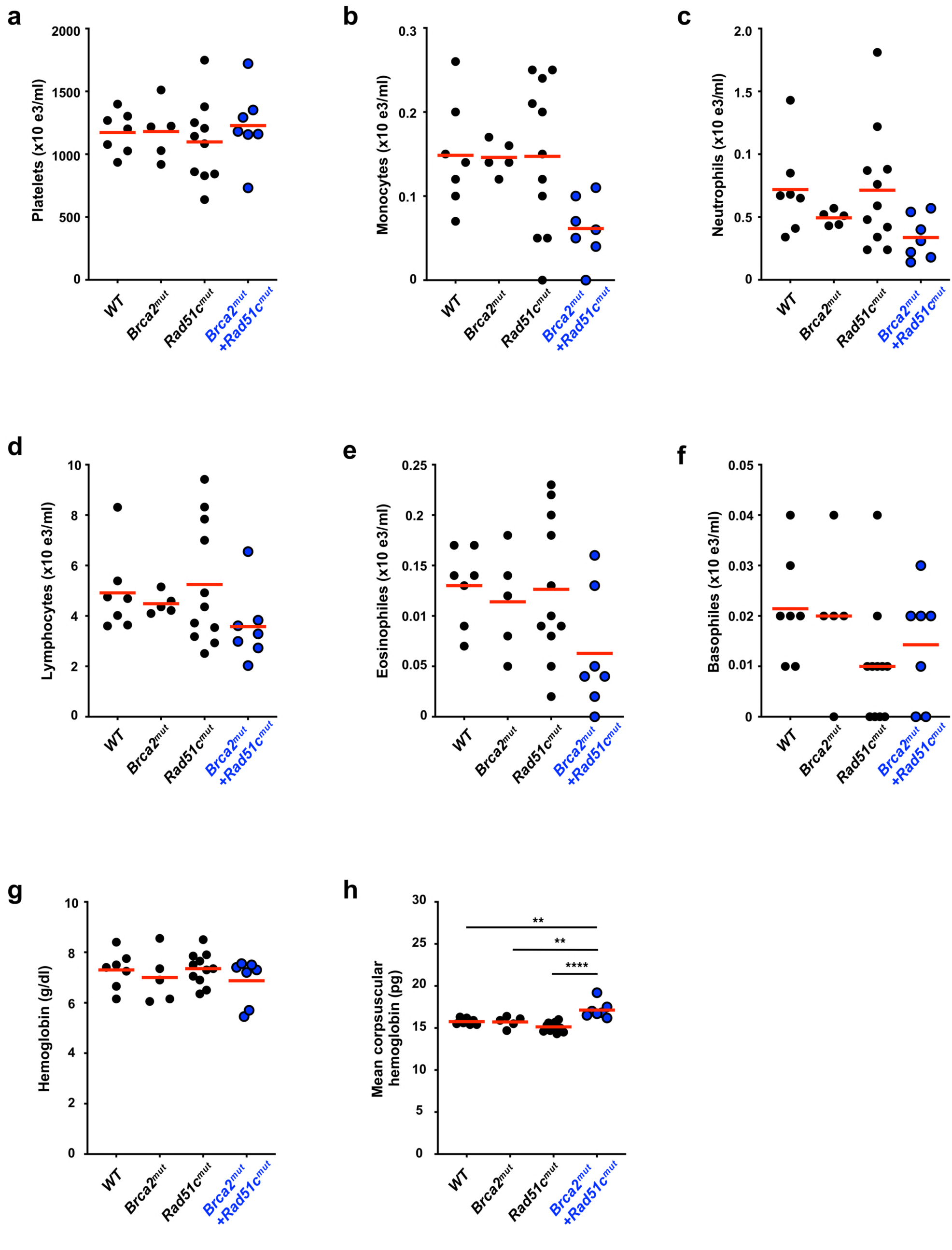
Complete blood count analysis of *Brca2*^*mut*^ *+Rad51c*^*mut*^ *mice*. **a-h**, Complete blood count analysis of 8–12 weeks old *Brca2*^*mut*^*+Rad51c*^*mut*^ and control mice (*n* = 6-12). **a**, Platelet count. **b**, Monocyte count. **c**, Neutrophils count. **d**, Lymphocytes count. **e**, Eosinophil count. **f**, Basophil count. **g**, Hemoglobin concentrations. **h**, Mean corpuscular hemoglobin concentration. Red bar in scatter graphs represent the mean. *p-*values are derived using the ANOVA test. ** *p* < 0.01, **** *p* < 0.0001.

**Extended Data Fig. 5.**
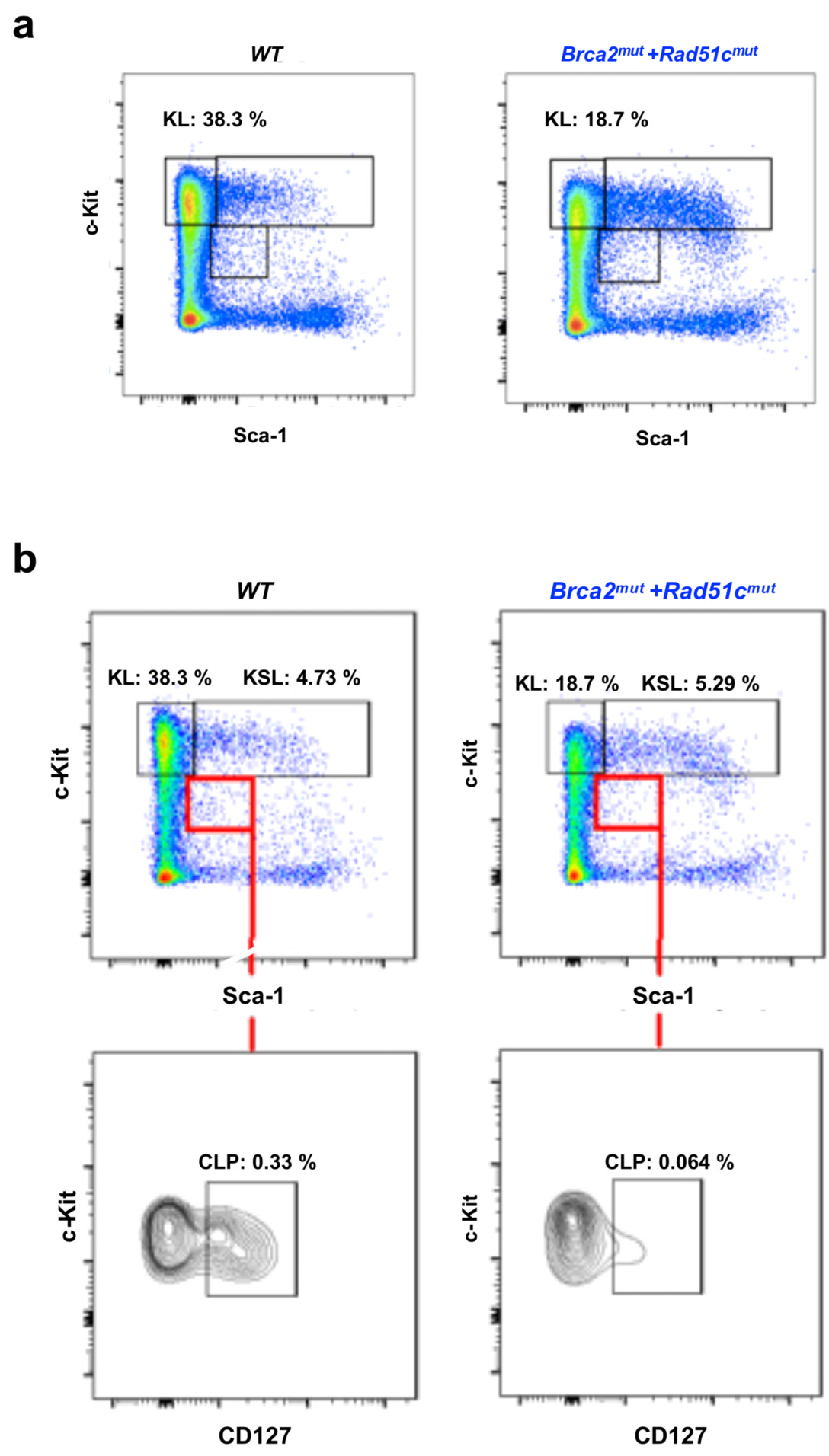
Hematological FACS analysis of *Brca2*^*mut*^*+Rad51c*^*mut*^ mice. **a**, Representative FACS plot showing gating for hematological lineage analysis performed for bone marrow cells collected from 8-12 weeks old mice with indicated genotypes (n=5-6). Lineage negative cells (20,000 events) are gated for CD117 (c-Kit) positive and Sca-1 negative populations to identify the KL subset designating myeloid progenitor cells. **b**, Representative FACS plot showing gating for identification of the CLP subset, designating common lymphoid progenitors. Lineage negative cells (20,000 events) are gated for c-Kit positive and Sca-1 moderate expression (KSL), which is further gated for CD127 positive cells to identify the CLP subpopulation designating the lymphoid population.

**Extended Data Fig. 6.**
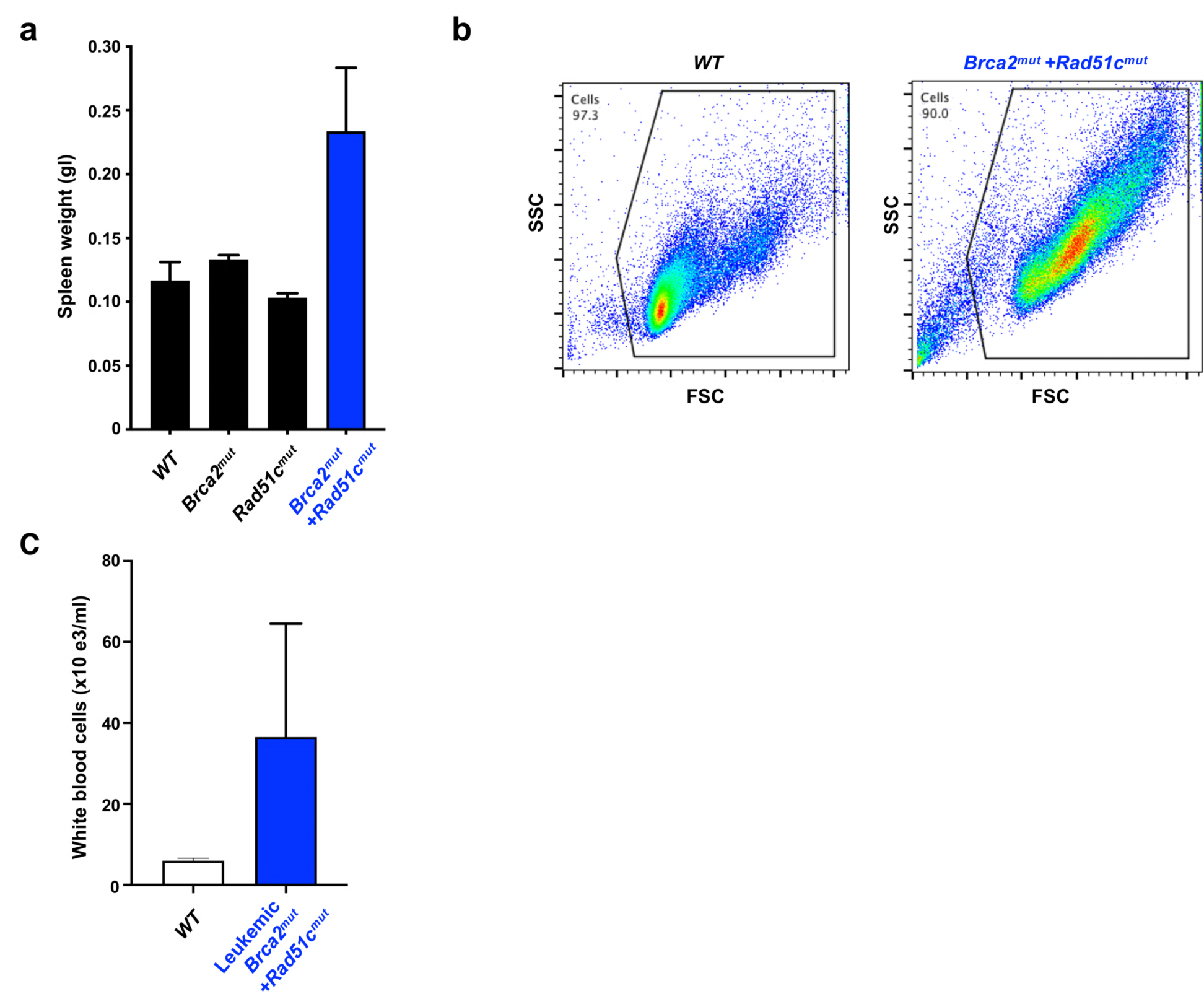
*Brca2*^*mut*^*+Rad51c*^*mut*^ mice show abnormalities in tissues of the immune system. **a**, Bar-graph of enlarged spleen quantification in malade *Brca2*^*mut*^*+Rad51c*^*mut*^ and age-matched control mice (n=3-6). **b**, Representative forward and side-ward scatter FACS plot image of thymus cells from WT and malade *Brca2*^*mut*^*+Rad51c*^*mut*^ mice show enlargement of the mutant cells, indicating T-cell activation. **c**, White blood cell count of leukemic *Brca2*^*mut*^*+Rad51c*^*mut*^ mice shows increased counts compared to age-matched wild-type (WT) mice (n=5-8). Error bars represent the standard error of the mean.

**Extended Data Table 1.** Polygenic *Brca2*^*mut*^*+Rad51c*^*mut*^ mice are born at sub-Mendelian frequency. Observed genotypes of *Brca2*^*mut*^*+Rad51c*^*mut*^ mice was determined by genotyping offspring of *Brca2* ^*mut/+*^*+Rad51c* ^*mut/+*^ intercross (total n=220). Expected genotypes are given using Mendelian genetics. *p*-value was calculated using chi-squared test (X^2^-test).

| Genotype | WT | *Brca2*<sup>mut/+</sup><br>*Rad51c*<sup>mut/+</sup> | *Brca2*<sup>mut</sup><br>+*Rad51c*<sup>mut</sup> | *p*-value |
| --- | --- | --- | --- | --- |
| Expected frequency | 6.25 | 25 | 6.25 | *p* = 0.02 |
| Observed frequency | 5.9 (n=13) | 29.5 (n=65) | 2.3 (n=5) |  |

**Extended Data Table 2.**
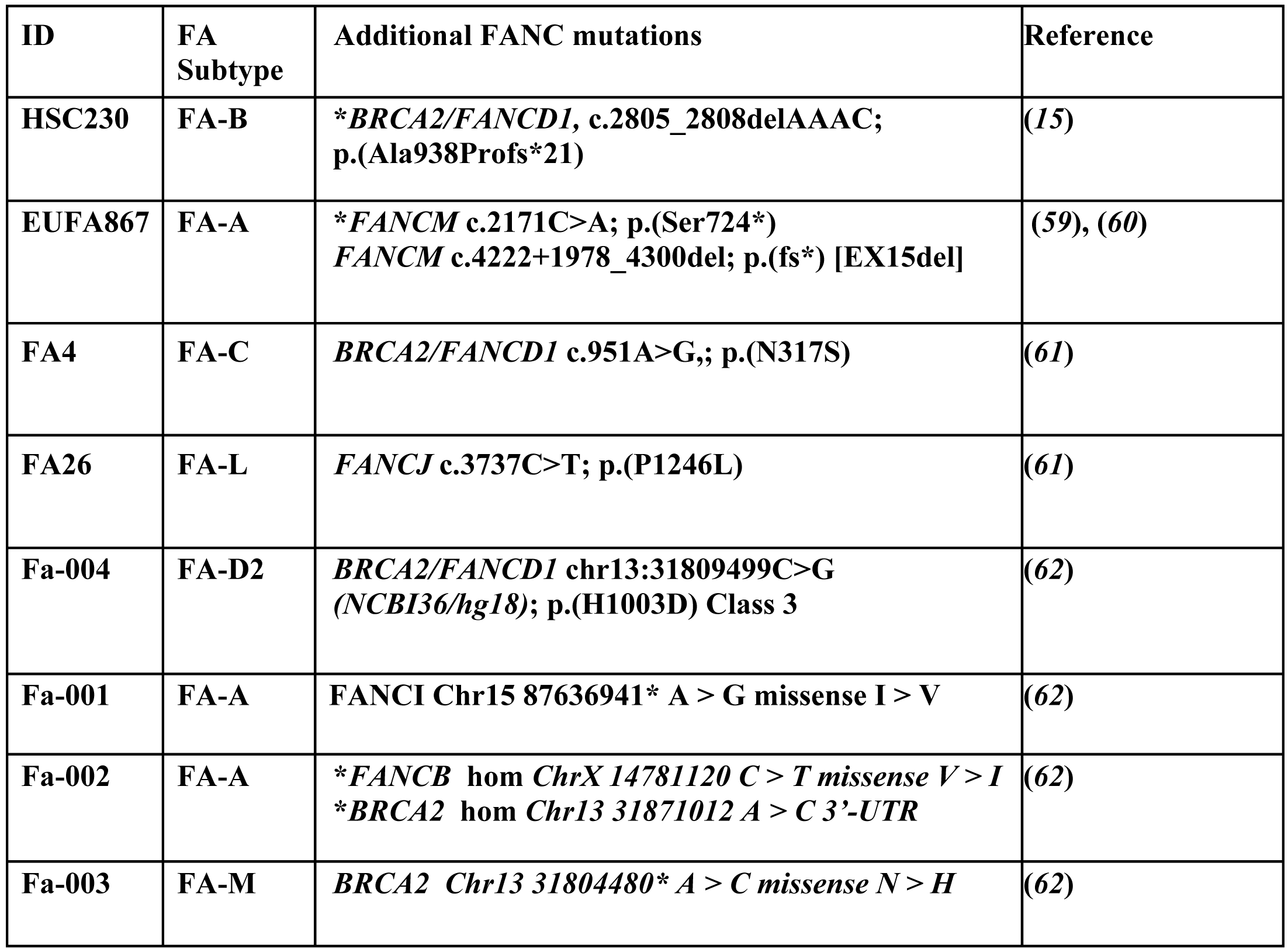
Previously reported *FANC* passenger mutations. Literature search of previously reported FA patients with germline *FANC* gene mutations in addition to a biallelic *FANC* inactivation. * denotes homozygous or biallelic inactivation of additional mutation.

